# Resolving spatiotemporal electrical signaling within the islet via CMOS microelectrode arrays

**DOI:** 10.1101/2023.10.24.563843

**Authors:** Anne Gresch, Jan D. Hüwel, Jennifer Briggs, Tim Berger, Ruben Koch, Thomas Deickert, Christian Beecks, Richard Benninger, Martina Düfer

## Abstract

Glucose-stimulated beta-cells synchronize calcium waves across the islet to recruit more beta-cells for insulin secretion. Compared to calcium dynamics, the formation and cell-to-cell propagation of electrical signals within the islet are poorly characterized. To determine factors that influence the propagation of electrical activity across the islet underlying calcium oscillations and beta-cell synchronization, we used high-resolution CMOS multielectrode arrays (MEA) to measure voltage changes associated with the membrane potential of individual cells within intact mouse islets. We measured both fast (milliseconds, spikes) and slow (seconds, waves) voltage changes and analyzed the spatiotemporal voltage dynamics. Treatment of islets from C57BL6 mice with increasing glucose concentrations revealed that single spike activity and wave signal velocity were both glucose-dependent. A repeated glucose stimulus involved a highly active subset of cells in terms of spike activity. When islets were pretreated for 72 hours with glucolipotoxic medium, the wave velocity was significantly reduced. Network analysis confirmed that the synchrony of islet cells was affected due to slower propagating electrical waves and not due to altered spike activity. In summary, this approach provided novel insight regarding the propagation of electrical activity and opens a wide field for further studies on signal transduction in the islet cell network.

**Article Highlights:** This study presents a new method for characterizing islet spatiotemporal electrical dynamics and subpopulations of beta-cells. We asked whether a high-resolution CMOS-MEA is suited to detect electrical signals on a level close to single cells, and whether we can track the propagation of electrical activity through the islet on a cellular scale. A highly active subpopulation of islet cells was identified by action potential-like spike activity, whereas slower waves were a measure for synchronized electrical activity. Further, propagating waves were slowed by glucolipotoxicity. The technique is a useful tool for exploring the pancreatic islet network in health and disease.

## Introduction

Glucose-induced depolarization of the beta-cell plasma membrane in the islet of Langerhans is a key property that underlies beta-cell function. Beta-cells are excitable and respond to nutrient stimuli, for example glucose. These nutrients, are metabolized, which drives an electrical response including membrane depolarization and elevated cytosolic calcium, which subsequently triggers exocytosis and insulin release (1).

Beta-cells in the islet form highly connected networks driven by connexin36 (Cx36) gap junction coupling (2,3). As a result oscillatory dynamics of membrane depolarization and subsequent changes in cytosolic calcium are synchronized, ensuring robustly glucose-regulated and pulsatile insulin release (3–5). Pancreatic beta-cells are functionally heterogeneous, and analysis of calcium dynamics has been used to classify leader, hub, or first-responder cell subsets (Figure 1A, 6). Imaging calcium dynamics is often used to infer spatiotemporal electrical dynamics (7–10). Conversely, direct recording of electrical activity, as for example by patch-clamping lacks spatial information.

**Figure 1:**
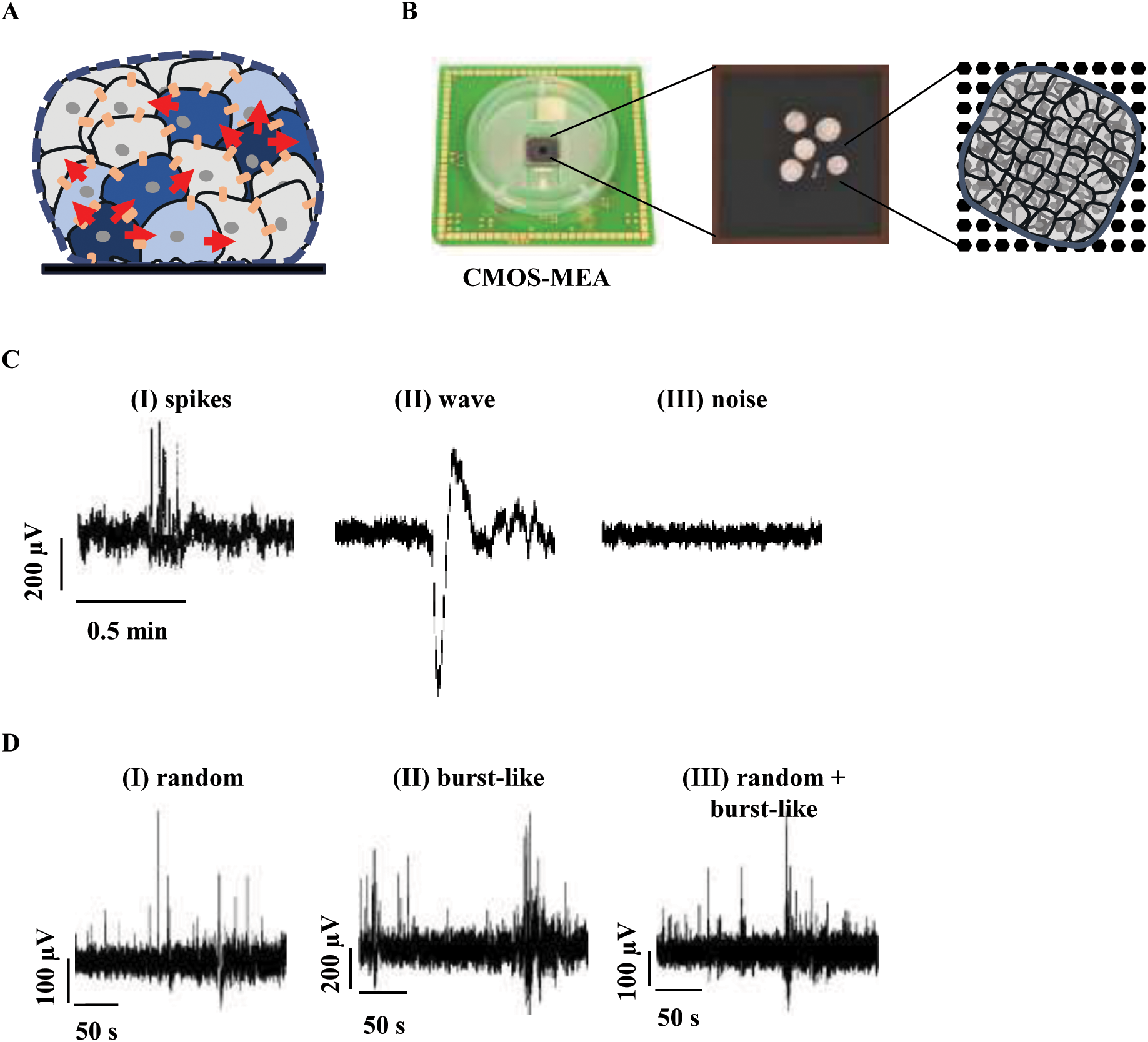
Signals of pancreatic islets measured with a high-resolution CMOS-MEA. A) Schematic representation of the beta-cell network in pancreatic islets, raising the possibility that synchronized oscillatory activity of cells is coordinated by a subset of beta-cells. B) CMOS-MEA chip with a culture chamber and zoom-in on the electrode array (black) with 5 islets applied. The second magnification shows a schematic representation of how one islet covers multiple electrodes. C) Three representative traces from three different electrodes with (I) AP-like signals, here referred to as spikes and (II) ion flux-driven waves. D) Three different electrode traces showing different forms of spike activity: (I) random, (II) burst-like or (III) a mixture of both.

An alternative method to study electrical dynamics relies on multielectrode arrays (MEAs) to measure the field potentials of cells within intact islets (11–14). An advantage of MEAs is that they provide a high temporal resolution (kHz) and are non-invasive. Patch-clamp or sharp electrode experiments are invasive and leak currents or channel-run down can affect membrane potential measurement and limits the recording duration. Previous MEA measurements (referred to as “conventional” MEAs) provided some limited spatial information with 1-3 electrodes per islet (14). However, this resolution is much less than provided by calcium imaging, which can resolve individual cells across intact islets.

Here we investigate the use of CMOS-MEA arrays with a high density of electrodes to record the spatiotemporal electrical dynamics across the islet. CMOS-MEA arrays have electrode sizes of 8 μm diameter, equivalent to the size of single islet cells. We examined whether CMOS-MEA arrays can resolve the spatiotemporal propagation and synchronization of electrical activity across the islet, and whether subsets of beta-cells with differing electrical activity or dynamics can be identified. We asked how important fast or slow time-scales of electrical activity are to these beta-cell subsets, and to the islet network as a whole. We further examined how glucolipotoxic conditions influence the spatiotemporal electrical dynamics.

## Research Design and Methods

### Islet preparation

Laboratory animal care was followed according to German laws (Az. 53.5.32.7.1/MS-12668, health and veterinary office Münster, Germany). Islets were isolated from adult male and female C57BL/6N mice (Charles River, Sulzfeld, Germany or in-house breeding, Münster, Germany). Mice were euthanized using CO_2_ and islets were isolated by collagenase digestion. After isolation, islets were kept in RPMI1640 medium (11.1mmol/L glucose) supplemented with 10% fetal calf serum, 100U/ml penicillin, and 100μg/ml streptomycin at 37°C in 5% CO_2_ humidified atmosphere.

### Electrical activity recording with microelectrodes array

1-2 days after isolation, islets were placed on the CMOS-MEA chips (Multi Channel Systems (MCS), Reutlingen, Germany) covered with 2 mL of culture medium. For experiments with glucolipotoxicity: 1 day after placing islets on the CMOS-MEA, culture medium was replaced by either control medium (0.28% fatty acid free BSA) or by glucolipotoxic medium (RPMI with 25mmol/L glucose and 100μmol/L sodium palmitate, 0.28% fatty acid free BSA) for 72 hours.

For measurements, the culture medium was replaced by 2mL bath solution. The CMOS-MEA was placed into the head stage, with heating applied from below and the array was covered with a dark chamber (as sensors are light sensitive). Electrical activity was initially examined before specific experimental protocols were conducted (Supplementary Material). Data acquisition was conducted with the CMOS-MEA-Control software (V2.8.0, MCS, Reutlingen, Germany). Data was filtered with a Butterworth 2nd Order filter between 0.1-500 Hz, processed at a frequency of 1 kHz and visualized using a Python script.

### Data Analysis

Our analysis uses two types of signals: one signal referred to as a *spike* (Figure 1C, trace I) and another signal as a *wave* with a negative and positive deflection (Figure 1C, trace II). Details on the signal extraction have been described previously (15,16 + Supplementary Material). Pearson-product-based network analysis was performed as previously described (8, 21). For each electrode pair within the area covered by the islet, the Pearson correlation coefficient was calculated using time series data over the indicated time window. The coefficients were computed to construct a correlation matrix (Figure 5A). An adjacency matrix was calculated by applying a threshold (0.5, 0.75 or 0.9) to the correlation matrix.

### Solutions and chemicals

Electrophysiological measurements were performed with a bath solution of (in mmol/L): 140 NaCl, 5 KCl, 1.2 MgCl_2_, 2.5 CaCl_2_, 10 HEPES, glucose as indicated, pH 7.4. Collagenase P was ordered from Roche Diagnostics (Mannheim, Germany), RPMI 1640, fetal calf serum and penicillin/streptomycin from Life Technologies (Darmstadt, Germany). BSA, diazoxide, nifedipine, tolbutamide, sodium palmitate from Sigma-Aldrich (Taufkirchen, Germany) or Diagonal (Münster, Germany).

### Statistics

Data was collected from islets of at least three independent mouse preparations for each series of experiments. Data are presented as scatter plots and/or means±SD. Gaussian distribution was determined by means of the Shapiro-Wilk test. For normally distributed data, statistical significance was assessed by unpaired Student’s t-test with Welch’s correction or by One-way-ANOVA followed by Tukey’s multiple comparison tests as indicated. For non-parametric distributions, statistical significance was assessed by Wilcoxon matched-pairs signed rank test, Friedman Test followed by Dunn’s for multiple comparisons or Kruskal-Wallis test followed by Dunn’s for multiple comparisons. GraphPad Prism Software (Version 9.5.1) was used for analysis. Values of p≤0.05 were considered statistically significant.

The separation between males and females did not yield a sufficient amount of data for all experiments to conduct reasonable statistic tests. Therefore, statistics comparing male and female islets were performed for selected data sets (Supplementary Material, Table 1).

## Results

### High-resolution detection of spatiotemporal electrical dynamics of islet cells in intact islets

The electrical activity of cells within intact islets was measured using a CMOS-MEA chip (Figure 1B, left), which incorporates 4225 electrodes and an inter-electrode distance of 16 µm. Several islets can be placed on each chip (Figure 1B, middle). For analysis, each islet was considered separately. Only the electrodes covered by the islet were included (Figure 1B, right), identified by exhibiting a stronger signal than the background noise (Figure 1C, compare III with I & II). Two different motifs were used for the analysis, one being very fast (∼millisecond) positively-oriented peaks similar to the shape of action potentials (APs, referred to as ‘spikes’, Figure 1C, I), and the other being pronounced waves consisting of a negative and a positive amplitude deflection (Figure 1C, II). Spikes occurred either randomly, in bursts or as a mixture of both patterns (Figure 1D). To compare the occurrence of spikes in measurements of different durations, we limited the spike detection to a duration of two minutes. When analyzing spike activity we did not distinguish between pattern of spikes, but only whether spikes were present or not.

### Glucose-triggered spikes depend on K_ATP_ and Ca_V_ channels

First, we examined how the voltage spike signals depended on glucose stimulation. Islets were sequentially stimulated with increasing glucose concentrations and activity was recorded 10 minutes after the start of each glucose stimulation. Figure 2A shows an example of CMOS-MEA electrodes (small squares) covered by an islet (grey squares) stimulated with increasing glucose concentrations, with electrodes showing spike activity highlighted in blue. The proportion of electrodes with spike activity, expressed as a percentage of all electrodes covered by an islet, showed a significant glucose dependence (Figure 2B). Next, we investigated ion channels that might be responsible for the spike signals and how spike activity was influenced by modulating membrane depolarization. In the presence of 10mmol/L glucose, the K_ATP_ channel opener diazoxide (250µmol/L) which hyperpolarizes the membrane (17), reduced spike activity to zero (Figure 2C). The L-type calcium channel blocker nifedipine (10µmol/L), also reduced spike activity, but not as markedly as diazoxide (Figure 2D). In the presence of 3mmol/L glucose, addition of the K_ATP_ channel blocker tolbutamide (100µmol/L) resulted in a significant increase in the recruitment of electrodes with spike activity (Figure 2E). Similarly, in the presence of 3mmol/L glucose, an elevated KCl concentration also increased the spike activity (Supplement Figure 1).

**Figure 2:**
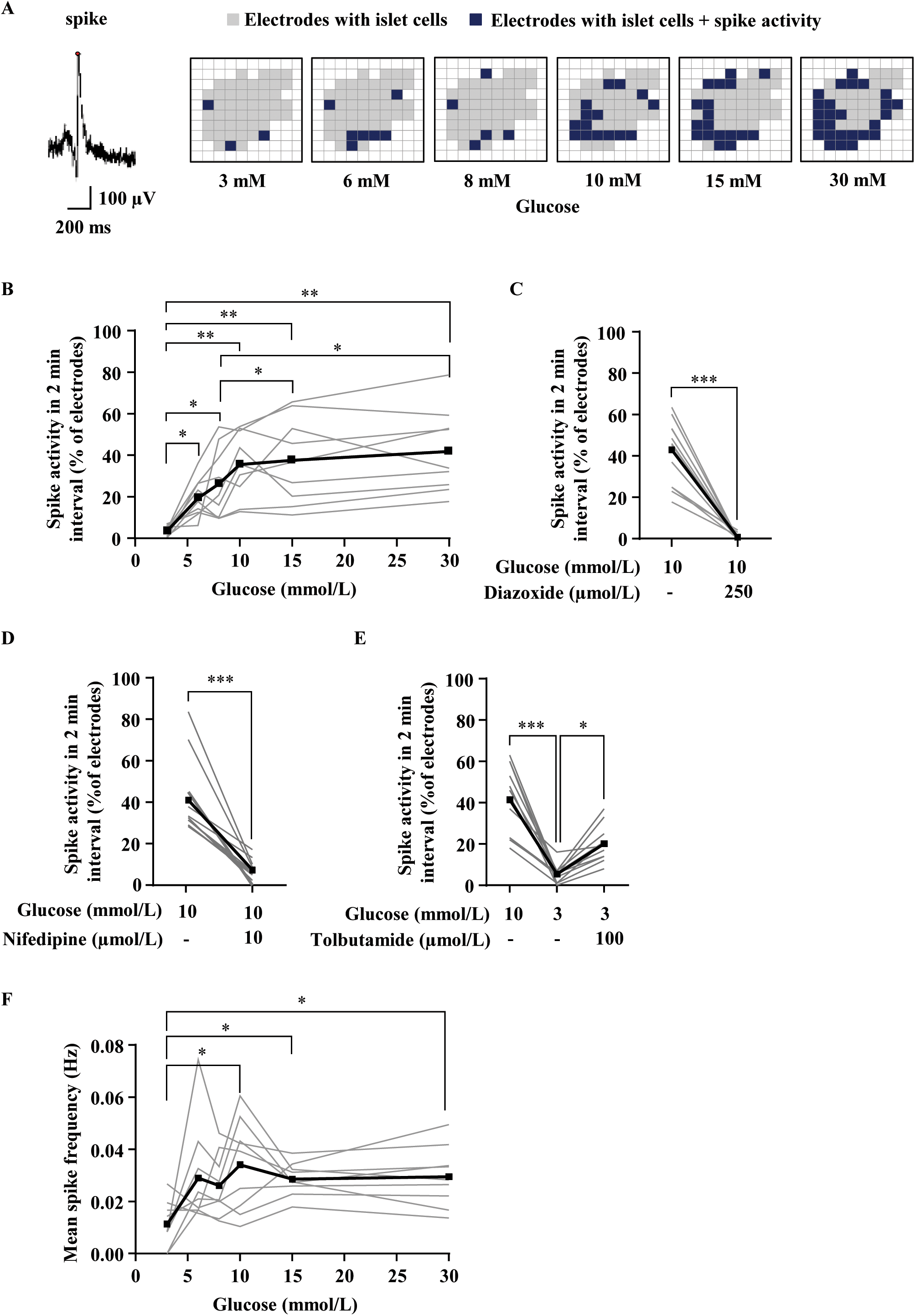
The occurrence of spike signals depends on glucose and can be altered by ion channel modulators. A) Image series of a representative islet showing which electrodes (small squares) are covered by the islet (gray) and which electrodes show spike activity (blue) within a 2-minute interval. B) The proportion of electrodes with spike activity expressed as a percentage of all electrodes covered by an islet shows a glucose dependence. C) The K_ATP_ channel opener diazoxide (250µmol/L) and D) the L-type calcium channel blocker nifedipine (10µmol/L) both significantly decrease spike activity, respectively. E) The K_ATP_ channel inhibitor tolbutamide (100µmol/L) increases the spike activity. F) The mean spike frequency of electrodes showing spikes increases with rising glucose concentration (same data set as in B). In B-E: The black thick line represents the mean value over all islets. Each gray line represents a single islet. In F: the gray line represents the average of the frequencies from all electrodes with spikes of one islet. Number of islets: 9 (B+F), 16 (C), 12 (D), 13 (E). Number of female/male mice from which islets were isolated: 1/4 (B+F), 1/3 (C), 2/2 (D), 2/1 (E). *p<0.05, **p<0.01, ***p<0.001.

To test whether spike dynamics themselves are altered by glucose, we evaluated the spike frequency, using the electrode data presented in Figure 2B. The mean spike frequency of those electrodes showing spikes rose with increased glucose concentration (Figure 2F), albeit with some variability. Similarly, the frequency increased with increasing glucose when considering all electrodes (including electrodes with no spikes) covered by the islet, with less variability (Supplement Figure 2A). Overall, these data show that increasing glucose recruits more cells displaying spikes and elevates the frequency of this electrical, AP-like signal.

### Repeated stimulus reveals consistent activity in a subset of islet cells

We next asked whether the same islet cells (electrodes) respond by showing spike signals upon a repetitive stimulus. Islets were stimulated three times in succession with 8mmol/L glucose. Between stimuli, the glucose concentration was reduced to 3mmol/L to bring cells to the same baseline level (Figure 3A, upper series of images). Activity was recorded 2 minutes after the addition of 8mmol/L glucose (to capture the first response) and 12 minutes after a change in the glucose concentration back to 3mmol/L. Consistent increases and decreases in spike activity were observed upon alternating between 3 and 8mmol/L glucose (Figure 3B). For the 3 stimuli, the number of times spikes were detected on an electrode was counted, and compared between all electrodes covered by an islet (Figure 3D, white bars). On average, 28±18% of the electrodes covered by islet cells showed spikes within the first minutes of stimulation. The percentage of electrodes, which always showed spikes (3/3 stimulations) is 11±9% under control conditions, suggestive of a population of highly active cells.

**Figure 3:**
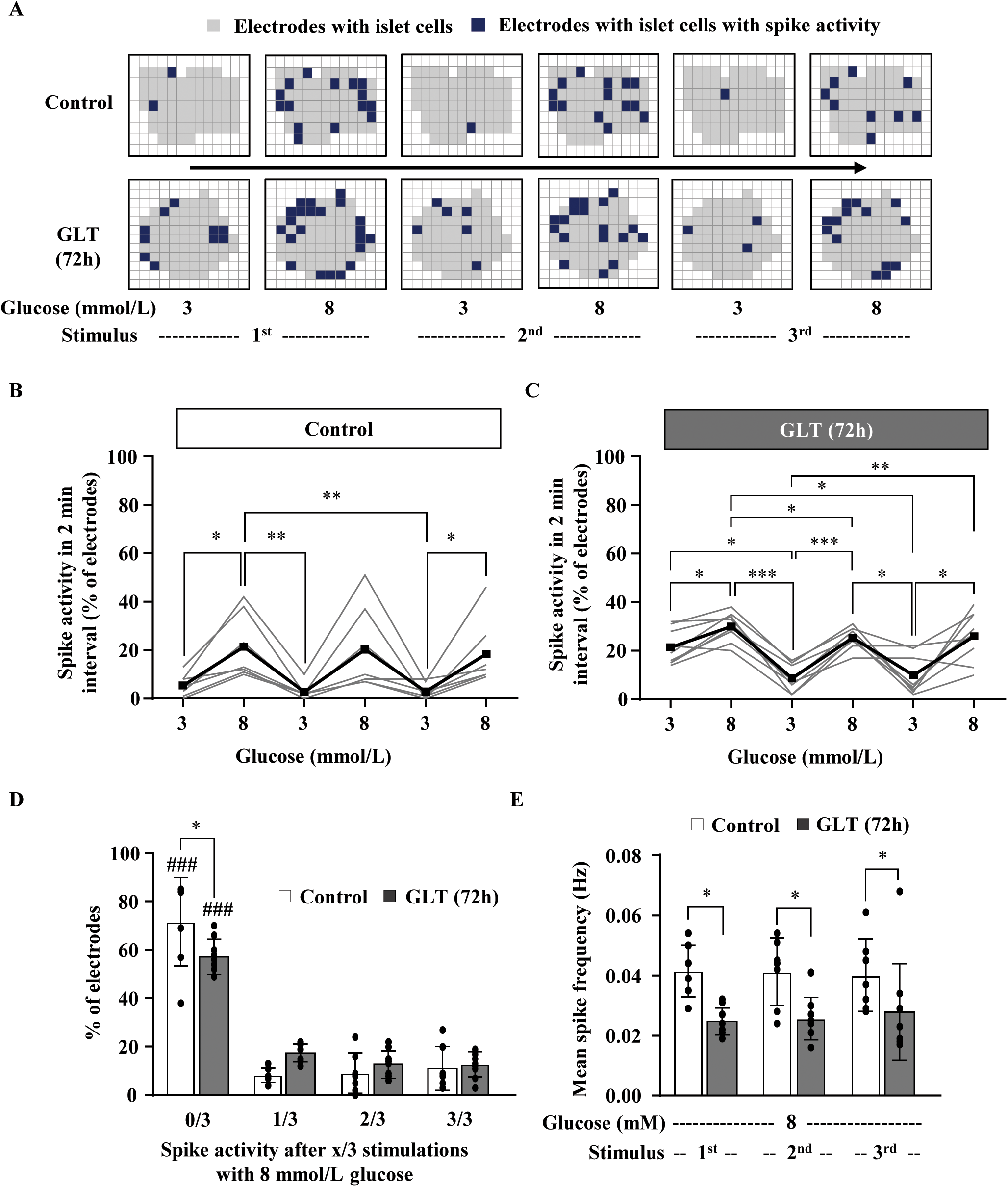
Glucolipotoxic conditions affect the recruitment of cells with spike activity. A) Image series of electrode array with one example islet for control (upper sequence of images) and one for GLT conditions (lower sequence of images). After 72 hours of pretreatment, islets were stimulated repeatedly (three times) with 8mmol/L glucose, with 12-minute breaks in between when glucose was reduced to 3mmol/L glucose. B) The proportion of electrodes with spike activity reaches each time the same level when stimulated with 8mmol/L glucose. C) Islets pretreated with GLT show a higher spike activity when a substimulatory glucose concentration of 3mmol/L is present, but still react as expected to 8mmol/L glucose. D) The number of times each electrode showed spike signals within the 3 stimuli was related to the total number of electrodes per islet. The islets treated with GLT medium show significantly fewer silent electrodes (in terms of spike activity) and tended to have more electrodes that showed one-time spike signals. E) The spike frequency of GLT-pretreated islets was lower in each case of stimulation with 8mmol/L glucose compared to control islets. Number of islets: control 7, GLT 9 (B-E). Number of female/male mice from which islets were isolated: 1/2 (B-E); *p<0.05, **p<0.01; ***p<0.001, ###p<0.001 *vs.* spike activity 1/3, 2/3, 3/3 Control and GLT (B).

Interestingly, when comparing the spike frequency of the 1^st^ and 2^nd^ phase of secretion, we observed a significant decrease when stimulated with 8mmol/L glucose (Figure 3E, 1^st^ phase 0.041±0.011Hz vs. Figure 2F, 2^nd^ phase 0.026±0.012Hz, p=0.012).

Islet cell dysfunction occurs in type 2 diabetes mellitus and glucolipotoxic conditions (18), therefore we tested if glucolipotoxicity affects the number of highly active cells, by culturing with medium supplemented with 25mmol/L glucose and 100µmol/L palmitate for 72 hours (GLT medium). For GLT-cultured islets, the number of electrodes that showed no spikes significantly decreased compared to control conditions (Figure 3D) and the number of electrodes that showed spike activity (1/3 times) tends to increase. Of note, the time at which this response appeared (i.e., at the first, second, or third stimulus) was variable (not shown). In contrast, the number of electrodes that showed highly active cells was similar between GLT and control conditions (Figure 3D, grey bar, 3/3 stimulations).

Compared to control conditions, the spike frequency in response to each stimulation with 8mmol/L glucose was significantly lower in GLT-treated islets (Figure 3E). In addition, islets exposed to GLT medium showed a higher percentage of electrodes with spike activity when glucose was initially lowered to 3mmol/L (Figure 3B and C, first 3mmol/L glucose treatment), which normalized in the following intervals (Figure 3C). The spike frequency in the presence of 3mmol/L glucose showed a trend to increase with GLT treatment, but it was not statistically different (Supplement Figure 2B).

Overall, data show that approximately 10% of the cells of an islet are highly responsive to repetitive glucose stimulation in terms of the spike signal. While more cells are responsive under glucolipotoxicity, their spike frequency is reduced.

### Membrane potential waves are increased by glucose stimulation and slowed by glucolipotoxicity

We next analyzed the waves in the electrical signal that were observed across the islet (Figure 4A, Supplement Figure 3). At 6-10mmol/L glucose, the wave duration was similar to that of fast calcium oscillations (3,19), in a range of ∼10 seconds, but decreased in response to 15mmol/L glucose (Figure 4A and B). To calculate the electrical wave velocity we used the information about the starting electrode and the absolute time difference of the signals identified by similarity analysis (Supplementary Material) between each electrode (Figure 4C). While the islet is a 3D structure, only a 2D plane is recorded by the chip as with most calcium imaging datasets. To account for potential variations in the velocity measurement by the distance or islet region used for the calculation, we chose 3 different distances from the wave start: 48µm (3 electrodes), 80µm (5 electrodes) and 160µm (10 electrodes). Similar wave velocities were measured for each distance used, suggesting uniform electrical wave propagation across the islet (Figure 4D). Thus, we used a distance of 80µm for subsequent wave velocity calculations. The velocity was variable, therefore only a trend in the increase of the velocity was observed upon increasing glucose concentration (Figure 4E). Nevertheless the velocity was in a similar range to that described previously for propagating calcium oscillations (70-150µm/s; 3,19). In contrast to the characteristics observed for calcium oscillations (3,19), we did not observe any correlation between islet size and electrical wave velocity (Figure 4F). However, the velocity was significantly reduced by the GLT treatment (Figure 4G).

**Figure 4:**
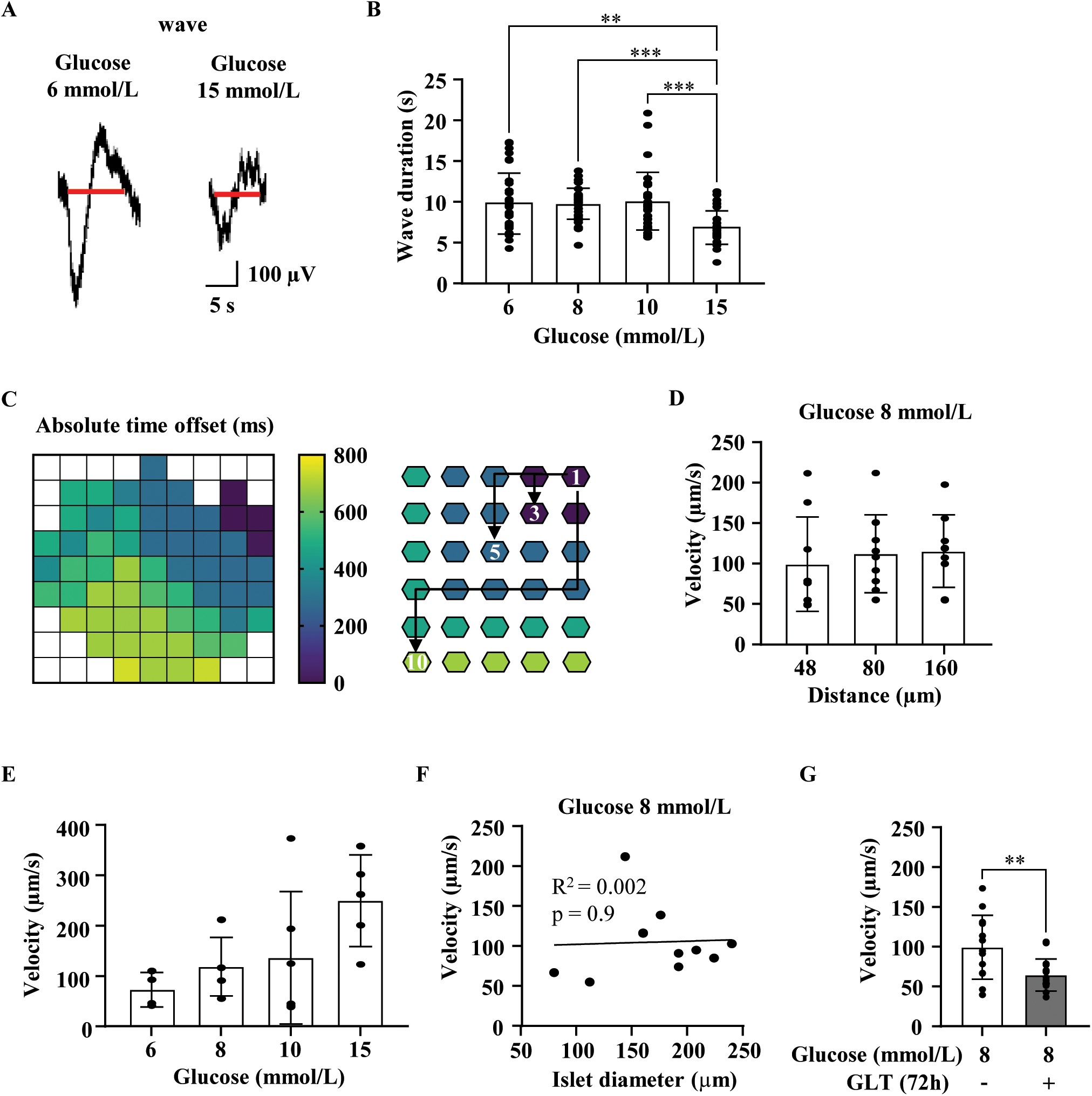
Spatiotemporal characterization of membrane potential waves propagating through the islet. A) Representative waves occurring with 6 or 15mmol/L glucose. The red line indicates how the wave duration was determined. B) Wave duration for different glucose concentrations decreases significantly with 15mmol/L glucose. Number of waves/islets: 32/9, 33/9, 31/6, 26/7. C) Heat map with absolute time offset in milliseconds of one islet treated with 8mmol/L glucose. On the right is a schematic of how the velocity per islet was determined over 3 different distances, i.e., the number of electrodes the motif passed through. D) The distance does not have any impact on velocity (n=9 islets). E) The wave velocity increased with elevated glucose concentrations (calculated over an 80 µm distance, n=4-6 islets). F) There is no correlation between islet size and velocity (n=11 islets). G) Pre-incubation with GLT medium significantly slowed down the wave propagation. Number of wave propagations/islets: 13/6, 16/6. Number of female/male mice from which islets were isolated: 1/3 (B, E), 2/3 (D), 2/4 (F), 1/4 (G). *p<0.05, **p<0.01; ***p<0.001.

### Intra-islet synchrony of waves increases with elevated glucose

Next, we examined whether the electrode time courses with respect to the wave and/or spike signals are synchronized, as is observed for calcium oscillations. For each pair of electrodes (covered by an islet), the Pearson correlation coefficient was determined. Figure 5A shows an example islet with the correlation coefficients at different glucose concentrations. The average correlation per islet increases with elevating glucose concentration, but is highly variable (Figure 5B). Interestingly the correlation at 10mmol/L glucose was less than that measured at 6-8mmol/L and 15mmol/L glucose, which is the glucose level where both spiking activity and spiking frequency peaked.

**Figure 5:**
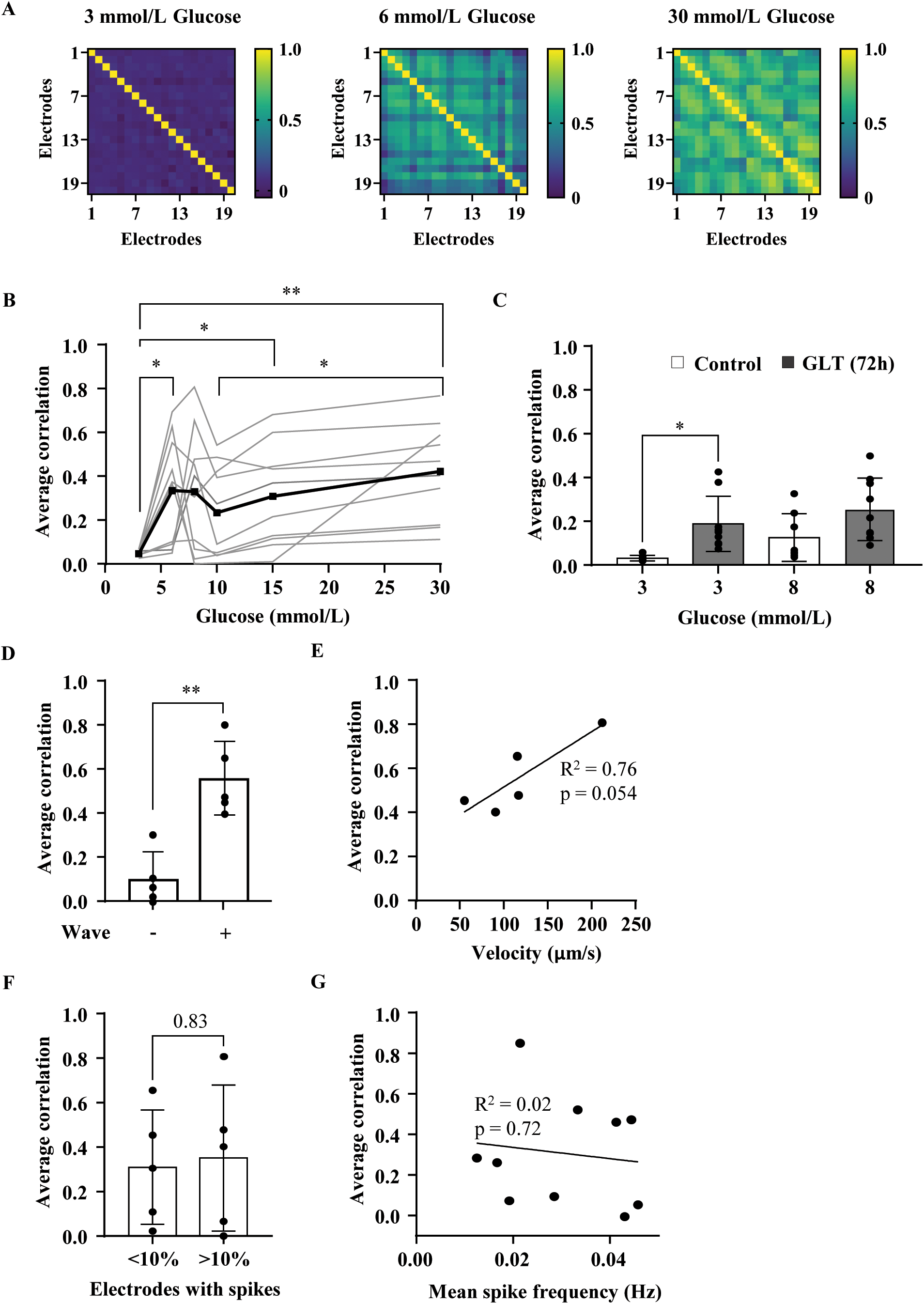
The average correlation depends mainly on the waves and not on the spike behavior of the cells. A) Representative heat maps of Pearson correlation coefficients for one islet treated with different glucose concentrations. B) Average correlation per islet depends on glucose concentration. C) Average correlation in the presence of 3mmol/L glucose is increased when islets are treated with GLT medium. D) The average correlation per islet is significantly higher when data set contains a wave motif. E) A high velocity contributes to an elevated average correlation. F) The average correlation factor seems independent of the amount of electrodes showing spikes involved. G) A slight negative relation exists between spike frequency and average correlation. Number of islets: 10 (B+D+F+G), control 7 /GLT 9 (C), 5 (E). Number of female/male mice from which islets were isolated: 1/4 (B, D, F, G), 1/2 (C, E). *p<0.05, **p<0.01.

We then analyzed whether the synchronization was still present when islets were pretreated with GLT medium (data from Figure 3C, 1st stimulus with 3 or 8mmol/L glucose). Surprisingly, islets pretreated with GLT medium showed higher mean correlation at 3mmol/L glucose (Figure 5C), but no difference at 8mmol/L glucose. When stimulating with 8mmol/L glucose, an increment was observed between the 1^st^ and 2^nd^ phases in terms of the average correlation (1^st^ phase, Figure 5C, control, 0.12±0.11 *vs*. 2^nd^ phase, Figure 5B, 8mmol/L glucose, 0.33±0.28, p=0.07).

We next examined the extent to which the synchronization depended on wave and spike features (using the data from Figure 2B, 8mmol/L glucose stimulus + 1 additional islet). If the time series data contained a propagating wave (5/10 islets), the mean correlation was significantly and substantially higher (Figure 5D). The mean correlation was also highly dependent on the velocity of the propagating wave (Figure 5E), with islets showing fast propagating waves exhibiting a high mean correlation. In contrast, the correlation was not affected by the spiking behavior: islets with less than 10% of electrodes (5/10 islets) showing spikes had a similar average correlation as islets with more than 10% of the electrodes showing spikes (Figure 5F). Note, although it is the same data set, the 5 islets that contain an electrical wave are not identical to the 5 islets where less than 10% of the electrodes showed spike activity.

Furthermore, there was no relation between the average correlation and spike frequency (Figure 5G). The average correlation for islets was similar if we considered only electrodes with spikes or without spikes (Supplement Figure 2C). These data suggest that the wave signal represents a marker or trigger for islet cell synchronization.

### Network properties of an islet derived from field potential measurement

Finally, to further examine the link between electrodes (cells) that are highly synchronized and slow waves, functional networks were constructed for each glucose concentration. Each electrode was considered as a node within a network (Figure 6A), with edges between nodes determined by a high correlation, above some threshold, which represents a functional connection. As shown previously with calcium imaging data (9), the functional connectivity within the islet is glucose-dependent, independent of the correlation threshold (Figure 6B). Network properties, including the clustering coefficient, global efficiency and local efficiency all followed the same pattern independent of the correlation threshold (Supplement Figure 4A-C). There was a significant correlation between cell connectivity and the wave duration (Figure 6C). Cells with the highest connectivity (normalized degree >0.6) tended to have longer wave duration (Figure 6D) and showed significantly greater voltage amplitude, in both positive and negative directions (Figure 6E and F). Thus cells showing the greatest synchronization also showed more robust longer waves.

**Figure 6:**
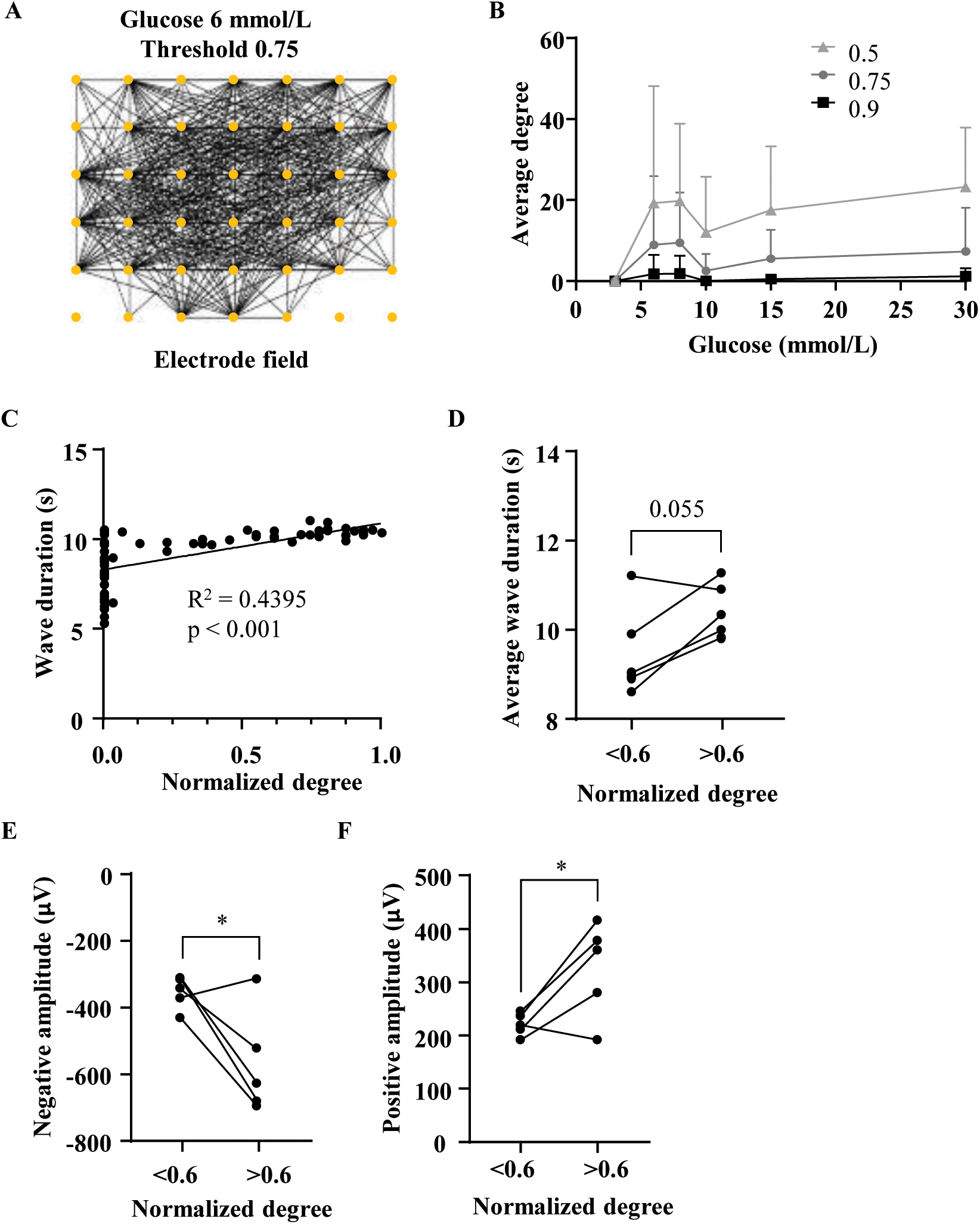
Islet cells with more functional connections show stronger wave signals. A) Visualization of islet network, with yellow dots representing electrodes and black lines are connections drawn when Pearson correlation coefficient was above set threshold (0.75). Glucose-dependent changes of the average degree (B) are calculated at different thresholds as indicated. C) Example islet where for each electrode the wave duration was correlated with the respective normalized degree. D) Summary of (C) for 5 islets showing a higher wave duration in highly connected cells. Additionally higher connected cells show a stronger deflection of the negative (E) and positive amplitude (F). Number of islets: 10 (B), 5 (D+E+F). Number of female/male mice from which islets were isolated: 1/4 (B, D-F). *p<0.05.

## Discussion

The CMOS-MEA enables recordings of the spatial-temporal electrical activity via measurement of the field potential across the islet. This approach can be used to characterize the spatiotemporal electrical response, heterogeneous responses throughout the islet, and breakdown under glucolipotoxic conditions. We analyzed two different voltage signals: *Spikes* which are AP-like signals generated by voltage-dependent calcium channels, and voltage *waves* resembling the characteristics of calcium oscillations. Both signals are glucose-dependent and show altered behavior in response to glucolipotoxicity. Through network analysis, we showed that the overall synchronization of islet cells depends on propagating waves rather than spike activity.

### High-speed multielectrode field potential recordings of the islet provides several advantages over other electrophysiological techniques

Typically when stimulated with >6mmol/L glucose the membrane potential of a mouse beta-cell depolarizes from -70 to -50mV from which APs start (1). Islet cells in our experiments exhibited spikes over an extended glucose concentration range (6 to 30mmol/L). This is a similar range (8.4 to 22mmol/L glucose) to mouse islets during patch-clamping (20). Prior work has used conventional MEAs where the whole islet covers 1-2 electrodes only (14,21). These prior studies also described two signals – single-cell APs and slow potentials – that are qualitatively similar to the fast spikes and waves respectively that we observe. For example, we observe a glucose dependence of the spike signal (Figure 2B, G) which is similar to the glucose-dependent APs previously described (14). Our data, suggests that the spikes result from voltage-dependent L-type calcium channels (Figure 2C-F), where L-type calcium channels drive mouse beta-cell APs (1). However, since nifedipine does not completely eliminate spike activity, other calcium channels such as R, N, or P/Q may also play a role (1). Overall, the CMOS-MEA measurements agree well with previous measurements and thus validate the recordings. However, a major advantage is that they provide additional spatial information at the level of individual cells.

Moreover, calcium imaging data has often restricted temporal resolution up to 4-8 Hz (3,22), which the CMOS-MEA chip far exceeds with a frame rate of 1 kHz (which can be extended to 25 kHz) while measuring a plane of cells. While the fast spike signals are likely due to calcium currents, field potential measurements have advantages over calcium imaging given that cytosolic calcium is influenced by transmembrane currents, as well as release from intracellular stores and export from the cytosol.

### CMOS-MEA recordings reveal a link between cellular electrical activity and islet synchrony

We observed a high level of synchrony or correlation between voltage time-courses of islet electrodes, which are qualitatively similar to the synchrony in calcium measurements between beta-cells that has been previously reported and depends on gap junction coupling (23). Our network analysis demonstrated that islets have synchronized electrical activity (Figure 5B). Importantly the slower electrical waves were associated with substantially higher correlated electrical activity (Figure 5D, E), which presumably travel through gap junctions across the islet. In contrast, the faster AP-like spikes do not contribute to the correlation of islet electrical activity (Figure 5F, G).

Functional analysis of calcium dynamics has classified subsets of beta-cells (6), such as hub cells which are defined as having more functional connections to other cells (7). We defined electrodes with more functional connections, which were associated with a much stronger wave signal (Figure 6C-F). This further supports a role for hub cells to provide greater electrical coordination across the islet. We also identified a subset of most active cells which always showed spike activity upon repeated glucose stimulation (Figure 3D). These cells made ∼11% of islet cells (electrodes). Whether these two subgroups (hubs and highly active) overlap remains to be clarified. However the lack of a link between AP-like spiking and synchronization suggests that these consistently spiking cells represent a different population or subset of cells. When the network analysis was performed with a series of glucose elevations, at 10mmol/L glucose, there was a small drop in the average correlation and for each network parameter (Figure 5B, 6B, Supplement 4A-C). The greatest variation in wave velocity was also observed at 10mmol/L glucose (Figure 4E), where wave velocity and average correlation are related (as discussed above, Figure 5E). At 10mmol/L glucose, islets reached a maximum spike frequency (Figure 2G) and these fast spikes lack any influence on the overall signal correlation (Figure 5G). Thus an increase in spike signal frequency generates an apparent decrease in correlation, whereas at further increases in glucose concentration are associated with increased velocity of slower waves that would provide an increase in correlation of electrical activity.

Previous work has suggested that biphasic insulin secretion consists of two electrical activity components: short trains of APs for the 1^st^ phase and a plateau depolarization interrupted by regular interburst intervals for the 2^nd^ phase (24). This suggests that fewer spikes will be found in the 2^nd^ phase, as we have actually observed (Figure 2F and 3E). In agreement with a prior study that used conventional MEA recordings (two electrodes per islet, 14), we found that synchrony within the islet increases from the 1^st^ to the 2^nd^ phase (Figure 5B and 5C). Our observation is also in agreement with calcium imaging data where the 1^st^ phase (activation phase) has a lower average node degree (equivalent to lower correlation) than the 2^nd^ phase (plateau phase) when stimulated with 8mmol/L glucose (8,9).

### A glucolipotoxic environment influences the spatiotemporal electrical dynamics of the islet

Glucolipotoxic environments have previously been found to have detrimental effects on beta cell function and cellular mass (18). A previous study used a conventional MEA and demonstrated via a single electrode recording of islet electrical activity that a glucolipotoxic environment increased the overall electrical activity of the islet (25). Consistent with this, we find that a glucolipotoxic environment increases the number of cells exhibiting spikes. The number of highly active cells was not altered by the glucolipotoxic conditions (Figure 3D), suggesting that our model pushes some cells to a lower glucose threshold, but this process is reversible upon the repetitive stimulation. We also observed that islets treated with glucolipotoxic conditions showed a lower spike frequency (Figure 3E), which is consistent with prior conventional MEA recordings that observed reduced AP frequency with glucotoxic conditions (14). These combined effects where single cell activity increased but with reduced spike frequency may reflect a compensatory mechanism; which enables present glucolipotoxic environments not being massively disruptive to insulin release (26).

It was previously shown that lipotoxic or glucolipotoxic milieus affect connectivity with the islet by reducing the proportion of functional links between cells (7), reducing Cx36 gap junction coupling (27) and disrupting calcium oscillation synchronization (28). Supporting the hypothesis of reduced coupling, we observed that the wave velocity decreased when islets were challenged by 72-hours glucolipotoxic environment (Figure 4G). In contrast our islets had a higher average correlation in electrical activity (Figure 5C) when exposed to glucolipotoxicity. However, this increment we observed at 3mmol/L glucose and may result from the lower glucose threshold increasing activity (Figure 5C).

In summary, we demonstrated that GLT conditions affected fast spiking by decreasing spike frequency, and slow wave electrical signals by slowing the propagation velocity. This likely does not have a major effect on intracellular calcium levels initially, given the additional influence of store-released Ca^2+^ release but could lead to a loss of coordination between cells over a longer term.

### Conclusion

In summary, we demonstrated that CMOS-MEA allows for high-resolution recording of spatiotemporal electrical activity across the pancreatic islet. We identified subsets of islet cells with a high response to glucose and which showed increased synchronization with the rest of the islet. The function of these cell subsets was not affected by a glucolipotoxic environment, but this environment affected the fast spike behavior of all cells and decreased signal propagation across the islet by decreasing connectivity between cells.

## Abbreviations

AP: action potential
CMOS: Complementary metal–oxide–semiconductor
Cx36: Connexin36
GLT: Glucolipotoxic(ity)
K_ATP_ channel: ATP-dependent potassium channel
kHz: kilohertz
MASS: Mueen’s algorithm for similarity search
MEA: multielectrode array

## Acknowledgement

We would like to thank Dr. Sven Schönecker for his continuous support on the CMOS-MEA device.

## Author Contribution

AG and MD designed the study. AG performed experiments of the study and analysed the data with the codes written by JDH, JB, TB, RK, TD and CB. AG wrote the manuscript, which was edited by AG, MD and RB. AG is the guarantor of this work and takes responsibility for the integrity of the data and the accuracy of the data analysis.

## Prior presentation information

Parts of this study were presented as poster presentation at the 83^rd^ Scientific Session of the American Diabetes Association in San Diego, California, USA, 23-26 June, 2023.

## References

1. Rorsman P, Ashcroft FM. Pancreatic β-Cell Electrical Activity and Insulin Secretion: Of Mice and Men. Physiological reviews 2018;98:117–214

2. Ravier MA, Güldenagel M, Charollais A, Gjinovci A, Caille D, Söhl G, Wollheim CB, Willecke K, Henquin J-C, Meda P. Loss of Connexin36 channels alters beta-cell coupling, islet synchronization of glucose-induced Ca2+ and insulin oscillations, and basal insulin release. Diabetes 2005;54:1798–1807

3. Benninger RKP, Zhang M, Head WS, Satin LS, Piston DW. Gap junction coupling and calcium waves in the pancreatic islet. Biophysical journal 2008;95:5048–5061

4. Head WS, Orseth ML, Nunemaker CS, Satin LS, Piston DW, Benninger RKP. Connexin-36 gap junctions regulate in vivo first- and second-phase insulin secretion dynamics and glucose tolerance in the conscious mouse. Diabetes 2012;61:1700–1707

5. Speier S, Gjinovci A, Charollais A, Meda P, Rupnik M. Cx36-mediated coupling reduces beta-cell heterogeneity, confines the stimulating glucose concentration range, and affects insulin release kinetics. Diabetes 2007;56:1078–1086

6. Benninger RKP, Kravets V. The physiological role of β-cell heterogeneity in pancreatic islet function. Nature reviews. Endocrinology 2022;18:9–22

7. Johnston NR, Mitchell RK, Haythorne E, Pessoa MP, Semplici F, Ferrer J, Piemonti L, Marchetti P, Bugliani M, Bosco D, Berishvili E, Duncanson P, Watkinson M, Broichhagen J, Trauner D, Rutter GA, Hodson DJ. Beta Cell Hubs Dictate Pancreatic Islet Responses to Glucose. Cell metabolism 2016;24:389–401

8. Stožer A, Gosak M, Dolenšek J, Perc M, Marhl M, Rupnik MS, Korošak D. Functional connectivity in islets of Langerhans from mouse pancreas tissue slices. PLoS computational biology 2013;9:e1002923

9. Stožer A, Skelin Klemen M, Gosak M, Križančić Bombek L, Pohorec V, Slak Rupnik M, Dolenšek J. Glucose-dependent activation, activity, and deactivation of beta cell networks in acute mouse pancreas tissue slices. American journal of physiology. Endocrinology and metabolism 2021;321:E305–E323

10. Dwulet JM, Briggs JK, Benninger RKP. Small subpopulations of β-cells do not drive islet oscillatory Ca2+ dynamics via gap junction communication. PLoS computational biology 2021;17:e1008948

11. Pfeiffer T, Kraushaar U, Düfer M, Schönecker S, Haspel D, Günther E, Drews G, Krippeit-Drews P. Rapid functional evaluation of beta-cells by extracellular recording of membrane potential oscillations with microelectrode arrays. Pflugers Archiv : European journal of physiology 2011;462:835–840

12. Schönecker S, Kraushaar U, Düfer M, Sahr A, Härdtner C, Guenther E, Walther R, Lendeckel U, Barthlen W, Krippeit-Drews P, Drews G. Long-term culture and functionality of pancreatic islets monitored using microelectrode arrays. Integrative biology : quantitative biosciences from nano to macro 2014;6:540–544

13. Raoux M, Bornat Y, Quotb A, Catargi B, Renaud S, Lang J. Non-invasive long-term and real-time analysis of endocrine cells on micro-electrode arrays. The Journal of Physiology 2012;590:1085–1091

14. Jaffredo M, Bertin E, Pirog A, Puginier E, Gaitan J, Oucherif S, Lebreton F, Bosco D, Catargi B, Cattaert D, Renaud S, Lang J, Raoux M. Dynamic Uni- and Multicellular Patterns Encode Biphasic Activity in Pancreatic Islets. Diabetes 2021;70:878–888

15. Hüwel JD, Gresch A, Berger T, Düfer M, Beecks C. Analysis of Extracellular Potential Recordings by High-Density Micro-electrode Arrays of Pancreatic Islets. In Database and Expert Systems Applications. Strauss C, Cuzzocrea A, Kotsis G, Tjoa AM, Khalil I, Eds. Cham, Springer International Publishing, 2022, p. 270–276

16. Hüwel JD, Gresch A, Koch R, Berns F, Düfer M, Beecks C. Tracing Patterns in Electrophysiological Time Series Data. In 2022 IEEE 9th International Conference on Data Science and Advanced Analytics (DSAA), IEEE, 2022, p. 1–10

17. Garrino MG, Plant TD, Henquin JC. Effects of putative activators of K+ channels in mouse pancreatic beta-cells. British journal of pharmacology 1989;98:957–965

18. Lytrivi M, Castell A-L, Poitout V, Cnop M. Recent Insights Into Mechanisms of β-Cell Lipo- and Glucolipotoxicity in Type 2 Diabetes. Journal of molecular biology 2020;432:1514–1534

19. Benninger RKP, Hutchens T, Head WS, McCaughey MJ, Zhang M, Le Marchand SJ, Satin LS, Piston DW. Intrinsic islet heterogeneity and gap junction coupling determine spatiotemporal Ca²⁺ wave dynamics. Biophysical journal 2014;107:2723–2733

20. Antunes C, Salgado A, Rosario Luis SR. Differential Patterns of Glucose-Induced Electrical Activity and Intracellular Calcium Responses in Single Mouse and Rat Pancreatic Islets. Diabetes 2000;49:2028–2038

21. Lebreton F, Pirog A, Belouah I, Bosco D, Berney T, Meda P, Bornat Y, Catargi B, Renaud S, Raoux M, Lang J. Slow potentials encode intercellular coupling and insulin demand in pancreatic beta cells. Diabetologia 2015;58:1291–1299

22. Langlhofer G, Kogel A, Schaefer M. Glucose-induced Ca2+i oscillations in β cells are composed of trains of spikes within a subplasmalemmal microdomain. Cell calcium 2021;99:102469

23. Briggs JK, Kravets V, Dwulet JM, Albers DJ, Benninger RKP. Beta-cell Metabolic Activity Rather than Gap Junction Structure Dictates Subpopulations in the Islet Functional Network, 2022

24. Misler S, Zhou Z, Dickey AS, Silva AM, Pressel DM, Barnett DW. Electrical activity and exocytotic correlates of biphasic insulin secretion from beta-cells of canine islets of Langerhans: contribution of tuning two modes of Ca2+ entry-dependent exocytosis to two modes of glucose-induced electrical activity. Channels (Austin, Tex.) 2009;3:181–193

25. Pajaziti B, Yosy K, Steinberg OV, Düfer M. FGF-23 protects cell function and viability in murine pancreatic islets challenged by glucolipotoxicity. Pflugers Archiv : European journal of physiology 2023;475:309–322

26. Schultheis J, Beckmann D, Mulac D, Müller L, Esselen M, Düfer M. Nrf2 Activation Protects Mouse Beta Cells from Glucolipotoxicity by Restoring Mitochondrial Function and Physiological Redox Balance. Oxidative medicine and cellular longevity 2019;2019:7518510

27. St Clair JR, Westacott MJ, Miranda J, Farnsworth NL, Kravets V, Schleicher WE, Dwulet JM, Levitt CH, Heintz A, Ludin NWF, Benninger RKP. Restoring connexin-36 function in diabetogenic environments precludes mouse and human islet dysfunction. The Journal of Physiology 2023;601:4053–4072

28. Hodson DJ, Mitchell RK, Bellomo EA, Sun G, Vinet L, Meda P, Li D, Li W-H, Bugliani M, Marchetti P, Bosco D, Piemonti L, Johnson P, Hughes SJ, Rutter GA. Lipotoxicity disrupts incretin-regulated human β cell connectivity. The Journal of clinical investigation 2013;123:4182–4194

