## Supplementary Material for "Resolving spatiotemporal electrical signaling within the islet via CMOS microelectrode arrays"

### Method Details

#### Initial examination of electrical activity

Electrical activity should be seen due a stimulatory glucose concentration of 10mmol/L. By changing from a high (10mmol/L) to a low (3mmol/L) glucose concentration, the electrical activity should disappear. This ensures the metabolic integrity of the islet under control conditions.

#### Signal extraction

The recorded data has the form of multiple time series of equal length, spatially arranged on a 65x65 grid. For evaluation, the grid was reduced to a rectangular area with the electrodes covered by an islet and some reference electrodes without islet cells.

*Spike detection:* For a time series $X = (x_1, \ldots, x_N)$, we first extracted all local maxima $\hat{X} = \{x_i \in X : x_{i-1} < x_i > x_{i+1}\}$ and then calculated the *peak score* (*PS*) for each of these points (S1). This measure is defined as $PS(X, i, k) = \frac{\sum_{j=i-k}^{i+k}(x_i - x_j)}{2k}$, where $x_i \in \hat{X}$ and $k$ defines the size of the interval used for comparison. The *PS* indicates how strongly a point exceeds its neighbors. Finally, the spikes in the time series are defined as the data points with significantly above-average *PS* that exceed a user-specified threshold (S2).

*Signal propagation tracker*: To examine the propagation of a wave motif, we developed the MoTrack algorithm (S3), which is based on the *MASS* algorithm (S4, S5) for calculating distances between signals. *MASS* is an algorithm for efficiently calculating the normalized Euclidean distance between a query signal and all subsequences of equal length in a time series. Due to the normalization, motifs that are structurally dissimilar can be sorted out, but motifs that only differ in amplitude can still be detected. MoTrack combines this approach with limits on temporal and spatial distances between waves to iteratively find occurrences of a propagated motif (S3).

#### Data Analysis

The wave crosses the baseline three times (Figure 4A). The *wave duration* was determined from the time between the first and last crossing of the baseline. The experimental data from Figure 2B+F was used for the analyses in Figure 4A-F, 5A-B, D-G, and 6.

**Table 1:** Comparison of female and male islet electrical activity in terms of number of electrodes with spikes, spike frequency, and wave propagation velocity.

| **Condition** | **Paraemeter** | **Male** | **Female** | **P-Value** |
|---|---|---|---|---|
| 10 mmol/L glucose | Spike activity in 2 min interval (% of electrodes) | 32±21, n=4 | 46±13, n=8 | 0.16 |
| | Mean spike frequency (Hz) | 0.04±0.01, n=4 | 0.04±0.01, n=8 | 0.88 |
| 3 mmol/L glucose | Spike activity in 2 min interval (% of electrodes) | 5±2, n=4 | 6±5, n=8 | 0.64 |
| | Mean spike frequency (Hz) | 0.021±0.009, n=4 | 0.04±0.03, n=8 | 0.3 |
| 3 mmol/L glucose + 100 µmol/L tolbutamide | Spike activity in 2 min interval (% of electrodes) | 20±2, n=4 | 20±11, n=8 | 0.92 |
| | Mean spike frequency (Hz) | 0.03±0.02, n=4 | 0.02±0.01, n=8 | 0.14 |
| 3 mmol/L glucose after 72 h GLT ($1^{st}$ stimulus) | Spike activity in 2 min interval (% of electrodes) | 25±7, n=5 | 16±2, n=4 | 0.04, * |
| | Mean spike frequency (Hz) | 0.01±0.02, n=5 | 0.025±0.005, n=4 | 0.12 |
| 8 mmol/L glucose after 72 h GLT ($1^{st}$ stimulus) | Spike activity in 2 min interval (% of electrodes) | 30±8, n=5 | 31±3, n=4 | 0.83 |
| | Mean spike frequency (Hz) | 0.027±0.006, n=5 | 0.023±0.001, n=4 | 0.24 |
| 8 mmol/L glucose | Velocity (µm/s) | 101±45, n=14 | 118±49, n=4 | 0.51 |

**Supplementary Figures**

**Figure 1**

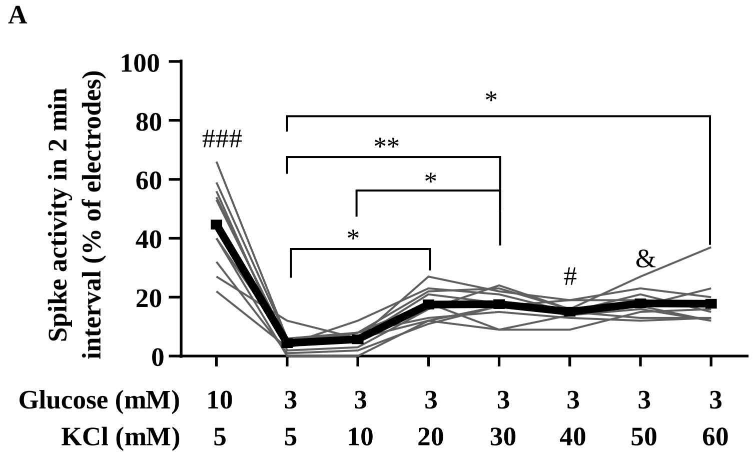


**Supplementary Figure 1:** The extracellular potassium concentration was successively elevated (with simultaneous reduction of sodium chloride to avoid hyperosmolarity) in the presence of a bath solution containing 3 mmol/L glucose. Each potassium (KCl) concentration was applied for 4 minutes before the next increase. The last two minutes were used for the evaluation in each case. An elevated extracellular potassium (KCl) concentration of 20 mmol/L (in the presence of 3 mmol/L glucose) resulted in a clear increase in the recruitment of electrodes with spike activity. A further increase of the KCl concentration (up to 60 mmol/L) only moderately augmented the number of electrodes involved. Number of female/male mice from which islets (n=10) were isolated: 2/2. *p<0.05, **p<0.01, #p<0.05 *vs.* 10 mmol/L glucose, &p<0.05 *vs.* 5 mmol/L and 10 mmol/L KCl, ###p<0.001 *vs.* 5 and 10 mmol/L KCl.

**Figure 2**

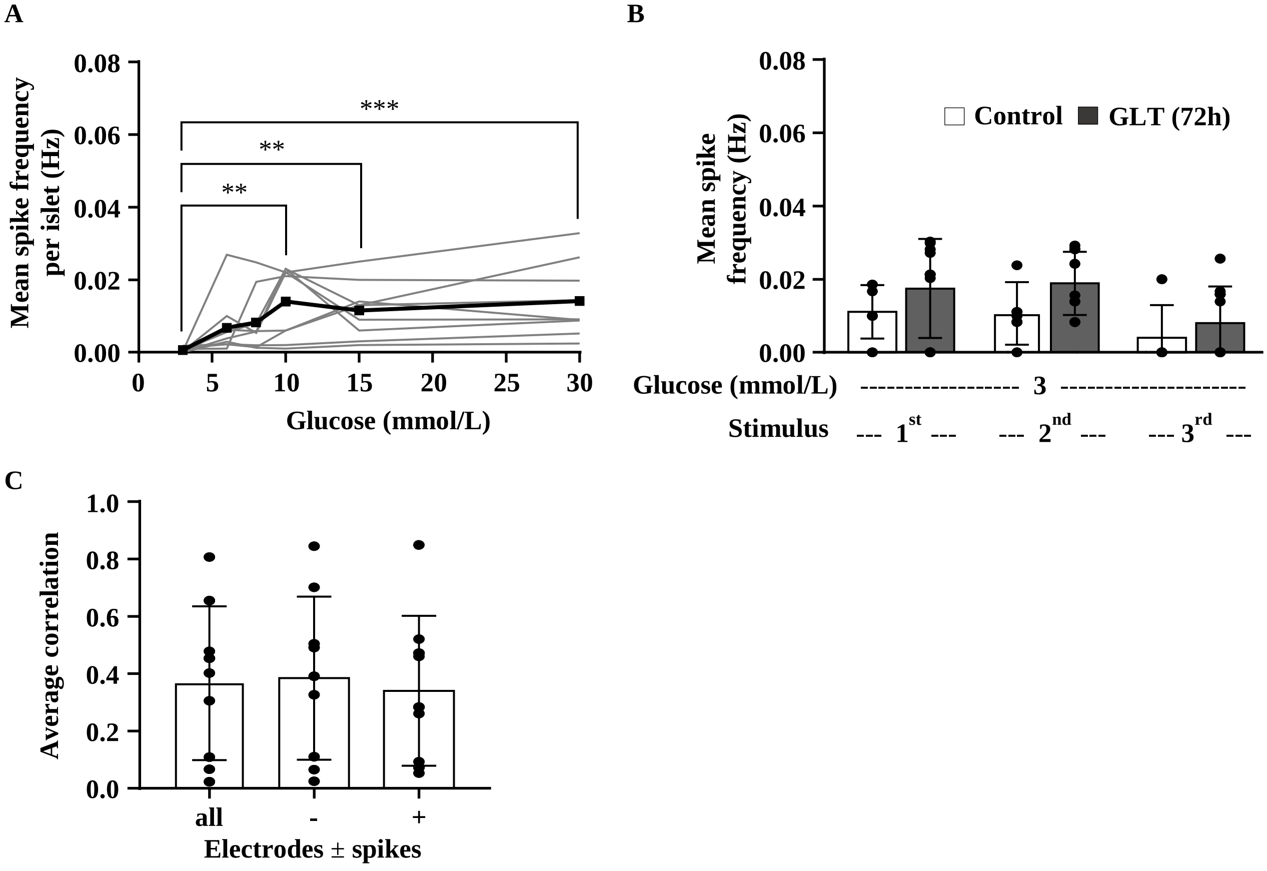


**Supplementary Figure 2**: A) The mean spike frequency was calculated over all electrodes covered by the islet independent of showing spikes or not. On average, the spike frequency per islet peaks at 10 mmol/L glucose. B) The mean spike frequency of electrodes showing spikes seems to be slightly elevated when islet are pre-treated with GLT medium. C) The average correlation over all electrodes of an islet, or just over the electrodes with or without spikes does not change. Therefore, spike signals do not influence the average Pearson-correlation-coefficient. Number of Islets: 9 (A), control 7 + GLT 9 (B), 9 (C). Number of female/male mice from which islets were isolated: 1/4 (A), 1/2 (B), 1/4 (C); **$p<0.01$, ***$p<0.001$.

**Figure 3**

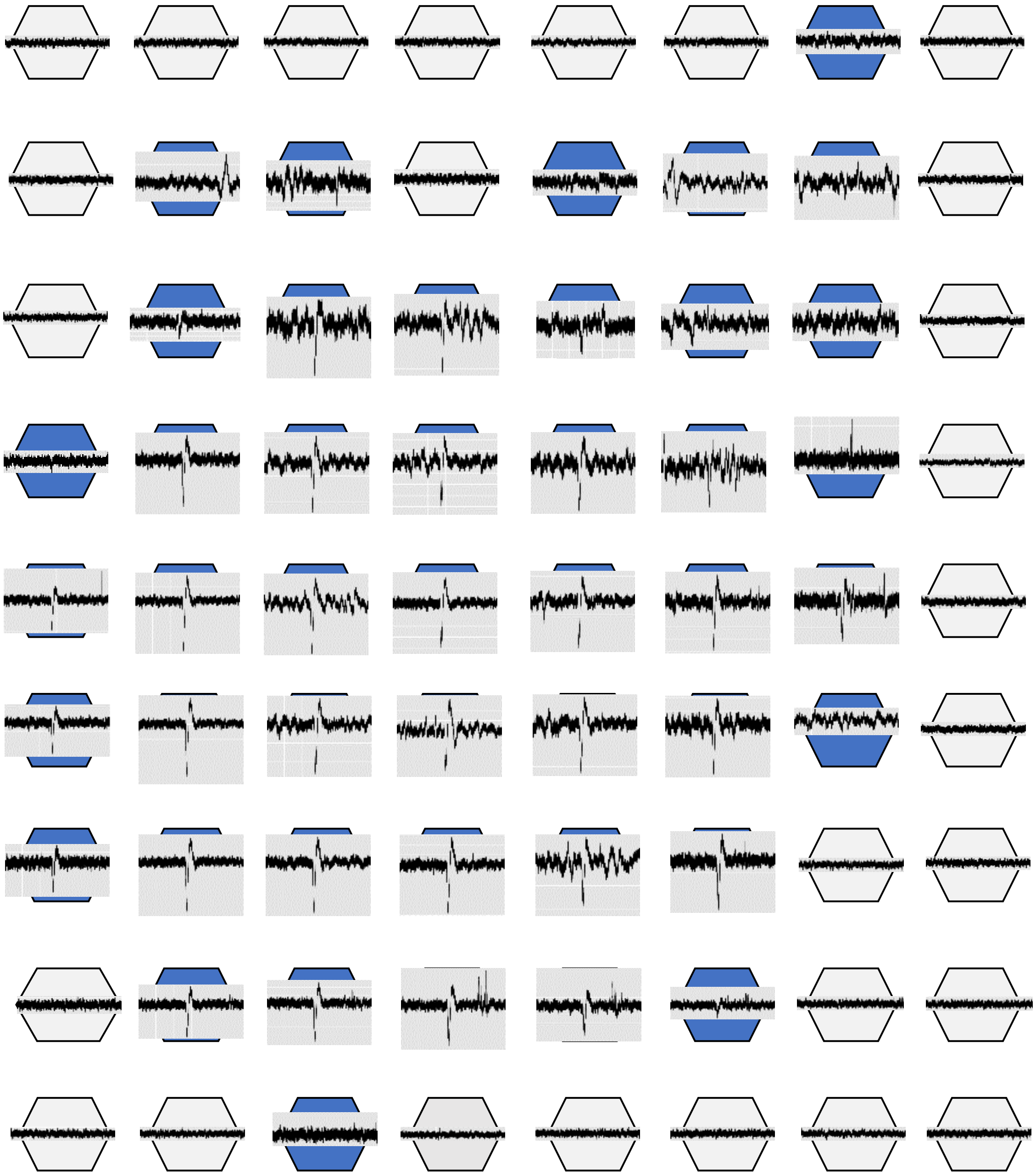

**Supplementary Figure 3**: Exemplary electrode field with voltage traces over a 2-minute interval, showing a wave signal after ~54 seconds in most of the electrodes covered by the islet (blue).

**Figure 4**

A

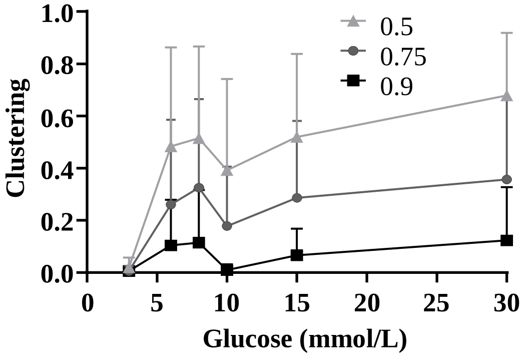


B

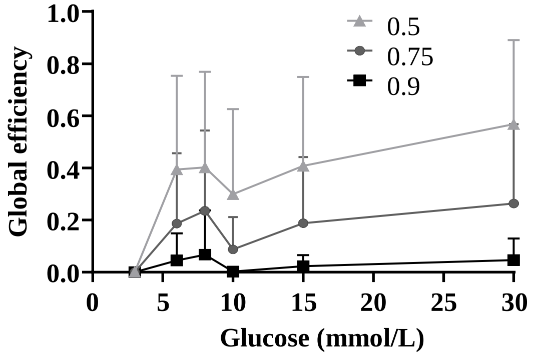


C

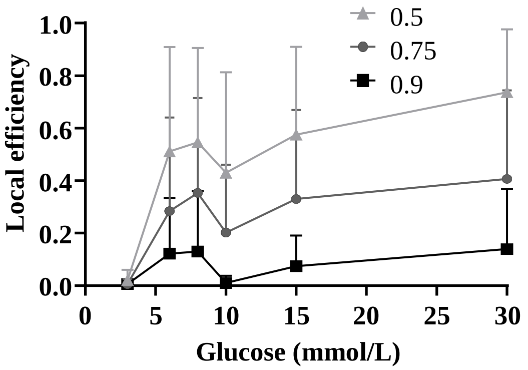


**Supplementary Figure 4:** Glucose-dependent changes in the average clustering coefficient (A), global (B) and local network efficiency (C) are calculated at different thresholds as indicated in the figures. Values of all parameters increase during glucose elevation but all show a small drop with 10 mmol/L glucose. Number of Islets: 10 (A-C). Number of female/male mice from which islets were isolated: 1/4 (A-C). No statistic was performed.

**Supplementary references**


S1. Palshikar GK. Simple Algorithms for Peak Detection in Time-Series. *Proceedings of 1st International Conference on Advanced Data Analysis* 2009; Business Analytics.

S2. Hüwel JD, Gresch A, Berger T, Düfer M, Beecks C. Analysis of Extracellular Potential Recordings by High-Density Micro-electrode Arrays of Pancreatic Islets. In *Database and Expert Systems Applications*. Strauss C, Cuzzocrea A, Kotsis G, Tjoa AM, Khalil I, Eds. Cham, Springer International Publishing, 2022, p. 270–276.

S3. Hüwel JD, Gresch A, Koch R, Berns F, Düfer M, Beecks C. Tracing Patterns in Electrophysiological Time Series Data. In *2022 IEEE 9th International Conference on Data Science and Advanced Analytics (DSAA)*, IEEE, 2022, p. 1–10.

S4. Mueen A, Zhong S, Zhu Y, Yeh M, Kamgar K, Viswanathan, Krishnamurthy, Gupta, Chetan Kumar, Keogh E. The Fastest Similarity Search Algorithm for Time Series Subsequences under Euclidean Distance. *URL: http://www.cs.unm.edu/~mueen/FastestSimilaritySearch.html.*

S5. Zhu Y, Imamura M, Nikovski D, Keogh E. Matrix Profile VII: Time Series Chains: A New Primitive for Time Series Data Mining (Best Student Paper Award). In *17th IEEE International Conference on Data Mining*. *18-21 November 2017, New Orleans, Louisiana: proceedings*. Piscataway, NJ, IEEE, 2017, p. 695–704.
